# Development of an efficient PEG-Mediated protoplast transformation system for the medicinal fungus *Ophiocordyceps xuefengensis*

**DOI:** 10.1101/2025.11.03.686195

**Authors:** Xiaoting Feng, Xinyao Sheng, Jun Liu, Rongrong Zhou, Zhongxu Yang, Xiaojuan Tang, Shuihan Zhang

## Abstract

*Ophiocordyceps xuefengensis* is a newly identified medicinal fungus with significant pharmacological and economic value, but its genetic manipulation has been impeded by the lack of an efficient transformation system. Here, we established the first stable and highly efficient polyethylene glycol (PEG)-mediated protoplast transformation platform for *O. xuefengensis* using hygromycin B as a selectable marker. Through systematic optimization of critical parameters such as enzyme composition, enzyme concentration, mycelial age, and digestion conditions, we developed an optimized protocol for protoplast preparation. A high protoplast yield of 9.42×10^7^ CFU/mL was achieved using 4-day-old mycelia with 1.5% lywallzyme 1 and 1.5% snailase digested at 34^◦^C with shaking at 130 rpm for 3.5 h. PY medium containing 0.6 M mannitol significantly enhanced protoplast regeneration. Stable plasmid integration and robust *hygR* gene expression were confirmed through PCR detection and sustained antibiotic resistance over four successive generations. Furthermore, a controllable expression system was established by using the endogenous promoter that drove the stable expression of the glycoside hydrolase gene *cbhI*. The result of enzymatic assay confirmed the functional production of CBHI enzyme, demonstrating the credibility of this system. This work provides a reliable genetic toolbox for functional genomics studies, targeted gene manipulation, and strain engineering in *O. xuefengensis*, facilitating fundamental research and sustainable utilization of this valuable medicinal species.

**IMPORTANCE:** The conservation and sustainable utilization of medicinal fungi represents a critical challenge in biotechnology and natural product research. *Ophiocordyceps xuefengensis*, a newly identified species with significant pharmacological potential, exemplifies this challenge: its wild resources are diminishing while artificial cultivation remains restricted by limited understanding of its fundamental biology. The lack of genetic tools has impeded progress in elucidating its biosynthetic pathways, regulating fruiting body development, and enhancing metabolite production. We established a highly efficient PEG-mediated transformation system that directly addresses this technological gap. In addition, we demonstrated that this transformation system can support heterologous gene expression using endogenous promoters. This platform enables reliable genetic manipulation in *O. xuefengensis*, permitting functional genomics studies and targeted bioactive compounds synthesis.

## INTRODUCTION

*Ophiocordyceps xuefengensis*, a new entomopathogenic fungus of the genus Cordyceps, discovered during the fourth national survey of the resources of Chinese materia medica (CMM) and recognized for its significant medicinal value [1]. As a sister taxon of *Ophiocordyceps sinensis* (*O. sinensis*) within the *Clavicipitaceae* family, it infects nymphs of insects *Phassus nodus* (Hepialidae) and develops a fruiting body on the surface of the host [2]. *O. xuefengensis* exhibits various pharmacological activities, including antibacterial, antiviral, and antitumor properties attributed to its rich content of bioactive compounds such as cordycepin, adenosine, polysaccharides, nucleotides, amino acids, and fatty acids [3, 4, 5, 6]. These properties underscore its potential for application in health products and medicine, positioning it as one of the most promising Cordyceps resources for further investigation.

Currently, active ingredients are primarily obtained from wild *O. xuefengensis*. However, sustainable utilization is challenged by the depletion of natural populations and the high cost of artificial cultivation, which is restricted by limited cultivation areas, low yields of bioactive components, and issues related to strain degeneration. Although a yeast cell factory for the production of ophiobolin-type sesterterpenoids has been successfully constructed in *Saccharomyces cerevisiae* [7], this approach remains unsuitable for industrial-scale production. Therefore, it is imperative to develop improved varieties with high active ingredient content and robust fruiting body yield to allow sustainable use of this valuable medicinal fungus.

Advances in genetic manipulation, transformation, and DNA integration techniques provide reliable strategies for the rapid development of transgenic medicinal fungi with targeted improvements in the size, biomass, and desirable agronomic traits of the fruiting body, as demonstrated in species such as *Cyclocybe aegerita* and *Cordyceps militaris* [8, 9]. Genetic transformation technology has become the preferred method for physiological and genetic research in fungi, as well as a key tool for generating strains with enhanced production of bioactive metabolites [10, 11]. Thus, the establishment of efficient genetic transformation technology presents a significant opportunity to accelerate the genetic improvement of *O. xuefengensis*.

The two primary methods for genetic transformation in filamentous fungi are polyethylene glycol-mediated protoplast transformation (PMT) and Agrobacterium-mediated transformation (AMT). Although AMT has been widely applied in monocots and cruciferous crops, it is often labor intensive, time-consuming, and exhibits low transformation efficiency in fungi [12, 13, 14]. Furthermore, the limited spore production of *O. xuefengensis* on artificial media makes AMT particularly challenging to implement. In contrast, PMT offers several advantages, including lower equipment requirements, simpler procedures, higher transformation efficiency, and stable inheritance of the transformed gene [15, 16, 17, 18]. This method relies primarily on the enzymatic removal of the fungal cell wall under optimized digestion conditions to generate protoplasts [19, 20], providing a more feasible and efficient approach for the genetic engineering of *O. xuefengensis*.

The composition of fungal cell walls shows a significant correlation between different species [21], making the selection of suitable hydrolytic enzymes critical to obtaining high-quality protoplasts in protoplast-mediated transformation studies [22]. Furthermore, transformation conditions profoundly influence the efficiency of DNA delivery, while the resuscitation and regeneration capacity of the protoplast directly determine the yield of transformants [23, 24]. To date, the PEG-mediated transformation method has been successfully applied and established several filamentous fungi, including *Cordyceps militaris* [25], *Aspergillus nidulans* [26], *Cordyceps javanica* [11], and *Pleurotus eryngii* [27].

However, the efficient PEG-mediated transformation (PMT) protocol has not yet be developed for *O. xuefengensis*, as an effective transformation method must be tailored to the biological characteristics of each fungal species. This limitation has impeded research on gene function, physiological and biochemical processes, and molecular mechanisms underlying fruiting body development in *O. xuefengensis*.

In this study, *O. xuefengensis* HACM001 was selected as experimental material, the protoplast preparation conditions, regeneration was systematically optimized and a simple, efficient and stable PEG-mediated protoplast transformation system of *O. xuefengensis* was successfully established. Furthermore, we applied this system to evaluate transformants via PEG-mediated random insertion. This represents the first successful attempt of direct genetic transformation in *O. xuefengensis* using PMT, demonstrating the feasibility of stable genetic manipulation in this species. This work provides a crucial technical platform for future genetic engineering and functional studies of this medicinal fungus.

## MATERIALS AND METHODS

### Strains and culture medium

The wild-type *O. xuefengensis* strain HACM001, was reserved in the Institute of Chinese Medicine Resources, Hunan Academy of Chinese Medicine, ChangSha, China [2]. The HACM001 strain was initially inoculated onto potato dextrose agar (PDA) medium at 28^◦^C in darkness for 10–14 days. Subsequently, a mycelial plug (0.5 × 0.5 cm) was aseptically excised from solid PDA medium using a flame-sterilized scalpel and transferred to 50 mL of PY liquid medium consisting of 15.0 g/L glucose, 5.0 g/L peptone, 2.0 g/L yeast extract powder, 1.0 g/L KH_2_PO_4_, 0.5 g/L MgSO_4_·7H_2_O. The culture was incubated at 25^◦^C with shaking at 130 rpm in the dark until adequate mycelial biomass was obtained for subsequent experiments.

### Preparation of protoplasts from *o. xuefengensis*

Mycelia were cultured in PY liquid medium and harvested by filtration through sterile gauze. The collected mycelia were thoroughly rinsed three times with 0.8 M mannitol and filtered again under the same conditions. Then 0.5 g of mycelia were gently mixed with 5 mL of the enzymatic lysis solution in the S1 buffer solution (0.8 M mannitol) and incubated at 30^◦^C . After incubation, the protoplast suspension was filtered and collected through four-layer filter paper and collected by centrifugation (4200 rpm, 10 min) at 4^◦^C. The pellet was washed twice with 10 mL of S2 buffer (0.6 M mannitol, 10 mM Tris-HCl pH 7.5, and 25 mM CaCl_2_). Finally, the protoplast pellets were resuspended with precooled S2 buffer and the protoplast yield was determined using a hemocytometer. All experiments were performed in triplicate and the data shown in the paper was the mean value.

### Optimization of protoplast preparation from *o. xuefengensis*

#### Influence of enzymatic systems and concentration on protoplast preparation

Six enzymatic lysis solutions combinations were prepared as follows: combination 1 consists of 1.5% lywallzyme 1 (Beijing Solarbio Science & Technology Co., Ltd); combination 2 consists of 1.5% lywallzyme 2 (Guangdong Institute of Microbiology, Guangdong, China); combination 3 consists of 50 U lyticase (Sigma-Aldrich Co., Ltd); combination 4 consists of 0.75% lywallzyme 1 and 0.75% snailase (Beijing Solarbio Science & Technology Co., Ltd); combination 5 consists of 0.75% lywallzyme 2 and 0.75% snailase, and combination 6 consists of 25 U lyticase and 0.75% snailase. Furthermore, the preparation of protoplasts was evaluated using combination 4 at different concentrations (0.75%, 1%, 1.25%,1.5%).

#### Influence of fungal age on protoplast preparation

Based on the optimized conditions established in Section 2.3.1, mycelia samples of *O. xuefengensis* were harvested in different cultures (4, 5, 6, and 7 days) and subjected to protoplast preparation according to the protocol described in Section 2.2.

#### Influence of enzymatic hydrolysis environments on protoplast preparation

The optimal protoplast yield of *O. xuefengensis* mycelia was investigated in varying enzymatic hydrolysis environments: time (2, 2.5, 3, 3.5, and 4 h), temperature (28, 30, 32, and 34^◦^C), and agitation speed (0, 50, 100, 130, and 150 rpm). The basic preparation protocol followed Section 2.2, with other conditions maintained as described in Section 2.3.2. The protoplast yields were quantified using a hemocytometer to determine the optimal enzymatic hydrolysis parameters.

### Screening for *o. xuefengensis* protoplasts regeneration medium

Three culture mediums (PPDA, PPY, and TB3) were evaluated for protoplast regeneration. A 200 µL aliquot of the *O. xuefengensis* protoplast suspension (1 × 10^7^ CFU/mL) was mixed with 5 mL of the respective liquid medium and incubated at 28^◦^ C overnight. After centrifugation, the pellet was resuspended in 400 - 600 µL of the corresponding medium and coated onto low melting point agar plates of the same composition. The plates were incubated at 28^◦^C in darkness for 6 - 7 days to assess the regeneration efficiency of the protoplasts. PPDA/PPY regeneration mediums were prepared by supplementing 0.6 M mannitol in PDA/PY medium, respectively. TB3 liquid medium consisted of 0.6 M mannitol, 3.0 g/L acid casein hydrolysate, 3.0 g/L yeast extract powder, 20.0 g/L sucrose.

### Selection pressure of *o. xuefengensis* to hygromycin B

The antibiotic sensitivity of *O. xuefengensis* strain HACM001 was determined using PDA medium supplemented with hygromycin B at six concentrations (100, 200, 300, 400, 500, 600 µg/mL). A control group without antibiotic supplementation was included. All petri dishes were incubated in darkness at 28^◦^C for 14 days.

### The polyethylene glycol-mediated transformation of *o. xuefengensis* protoplast

A 200 µL aliquot of protoplast suspension was transferred to a sterile 50 mL centrifuge tube and gently mixed with 6 µg of the linearized plasmid pCAMBIA1300 carring hygromycin B resistance (*hygR*) gene. The mixture was incubated on ice for 5 min. Then 50 µL of S3 buffer (25% PEG4000, 10 mM Tirs-HCl pH 7.5, and 25 mM CaCl_2_) was added dropwise to the mixture with gentle agitation. After an additional 30 min of incubation on ice, 1 mL of S3 buffer was carefully introduced along the inner wall of the tube, followed by incubation at room temperature for 5 min. The mixture was then transferred to 5 mL of liquid PPY regeneration medium and incubated overnight at 28^◦^C with shaking at 100 rpm. After centrifugation, the pellet was resuspended in PPY liquid medium and transferred to 30 mL of molten PPY low melting point medium containing 600 µg/mL hygromycin B for plate pouring. The plates were incubated in the dark at 28^◦^C for 6-7 days. Putative transformants that appeared after four rounds of screening in antibiotic-containing medium were considered positive transformants.

### Transformant Verification

Total genomic DNA extracted from both wild-type O. xuefengensis an-d positive transformant strains was used as a template for PCR amplification with the primer pair (PF1: CTTATATGCTCAACACATGAGCG; PR1: ATCTCCACTGACGTAAGGGATGAC).

The amplification products were then analyzed by agarose gel electrophoresis to determine their molecular weights.

### PEG-Mediated Protoplast Transformation of the pCAMBIA1300-CBHI Expression Cassette

Cellobiohydrolase (CBHI) is the main cellulase. The genes of CBHI (PDE_07945) from *penicillium oxalicum* 114, endogenous *gpd1* promoter (p*gpd1*) from *O. xuefengensis*, together with the terminator of *trpc* from *Aspergillus niger*, were subcloned into pCAMBIA1300 vector backbone to generate the recombinant plasmid pCAMBIA1300-CBHI, which was subsequently transformed into *O. xuefengensis* host cells.

### Validation of the CBHI Expression and Vitro Enzyme Activity

The positive recombinant *O. xuefengensis*-CBHI transformants were randomly selected by selecting single well-growing colonies from medium containing 600 µg/mL hygromycin B. These were transferred to fresh PY medium with the same antibiotic concentration and cultured in darkness at 28^◦^C for 7 days. The mycelium samples were then harvested for genomic DNA extraction, the DNA obtained was used as a template for target gene amplification, following the experimental procedures outlined in Section 2.7 with the primer pair (PF2: GGTGGCTACCTGAGCAGGGA; PR2: ACGGCTGCACTGAACGTCAG).

For CBHI production, two positively recombinant transformants were cultured on resistance plates and then cultured in a 500 mL triangular flask containing 100 mL of PY liquid medium at 30^◦^C with shaking at 130 rpm for 96 h. After fermentation incubation, extracellular protein was collected from the culture supernatant and purified using His TrapTMFF column. The CBH activity was measured according to the method described by Song et al. [28].

## RESULTS

### The preparation of *O. xuefengensis* protoplasts under different factors

#### Effect of enzymatic systems and concentration on protoplasts preparation

The effect of the preparation of fungal protoplasts varies due to the difference in the structures of the fungi’s cell wall, and specific enzymes were required to remove the cell wall during the protoplast preparation process [29, 30]. Based on previous studies [31, 32], six different enzyme combinations were selected for enzymatic digestion of *O. xuefengensis* mycelia. Protoplast production varied significantly between different enzymatic treatments (Fig. 1A). Notably, the dual-enzyme systems consistently outperformed single-enzyme preparations in terms of protoplast production. Among binary enzymatic combinations, combination 4 consists of 1.5% lywallzyme1 and 1.5% snailase produce the highest protoplasts with a concentration of 2.23 × 10^6^ CFU/mL, followed by combination 5 (1.46 × 10^6^ CFU/mL), and combination 6 resulted in the lowest yield (1.25 × 10^6^ CFU/mL), with all differences statistically significant (p < 0.01). For a single enzyme, a yield of 1.05× 10^6^ CFU/mL produced by 1.75% lywallzyme 2 alone, which was not statistically significant difference (p > 0.05) from the yield obtained treatment with 1.5% lywallzyme 1 (0.88 × 10^6^ CFU/mL) and 50 U lyticase (0.85× 10^6^ CFU/mL). Therefore, combination 4 was selected as the preferred enzyme for the protoplast preparation from *O. xuefengensis*.

**FIG 1.**
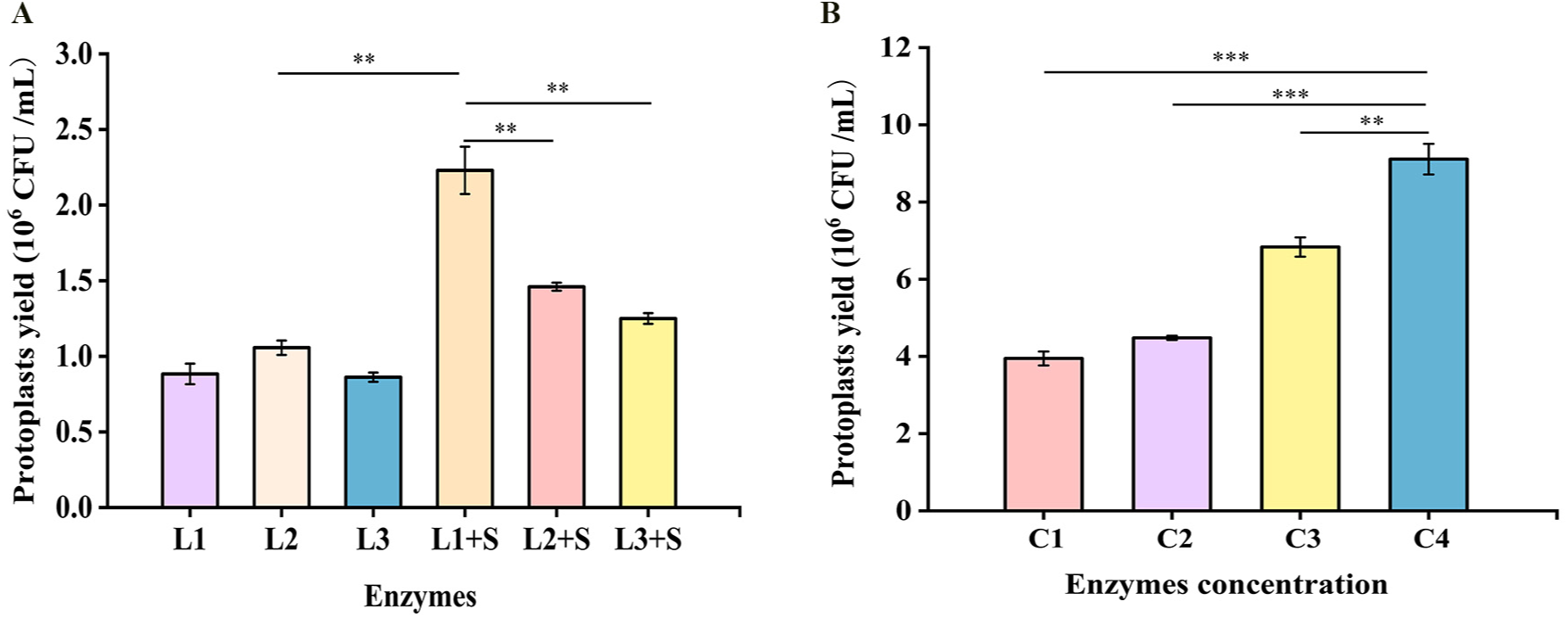
Yield of *O. xuefengensis* protoplasts of different enzyme and concentration. (A) Types of enzymes: L1: 1.5% lywallzyme 1, L2: 1.5% lywallzyme 2, L3: 50 U lyticase, L1 + S: 0.75% lywallzyme 1 and 0.75% snailase, L2 + S: 0.75% lywallzyme 2 and 0.75% snailase, L3 + S: 25 U lyticase and 0.75% snailase; (B) enzyme concentration: C1: 0.75% lywallzyme 1 and 0.75% snailase, C2: 1% lywallzyme 1 and 1% snailase, C3: 1.25% lywallzyme 1 and 1.25% snailase, C4: 1.5% lywallzyme 1 and 1.5% snailase. Values are presented as mean ± SD (*n* = 3). Significance analysis between two different groups:^∗^ *p <* 0.05, ^∗∗^ *p <* 0.01, and ^∗∗∗^ *p <* 0.001.

In addition, this study investigated the effect of enzyme concentration on protoplast preparation. A significant positive correlation was observed between protoplast yield and enzyme concentration, indicating the adaptability of *O. xuefengensis* to varying enzymatic conditions during protoplast isolation (Fig. 1B). The highest protoplast yield was achieved at an enzyme concentration of 1.5%, reaching 9.11 × 10^6^ CFU/mL. This value was markedly higher than protoplast production at an enzyme concentration of 1.25% (3.95 × 10^6^ CFU/mL, p < 0.01), 1% (4.48 × 10^6^ CFU/mL, p < 0.001), and 0.75% (5.83 × 10^6^ CFU/mL, p < 0.001). These results underscore the critical importance of optimizing enzyme concentration to maximize protoplast yield during enzymatic digestion.

#### Effect of fungal age on protoplast preparation

The preparation of protoplasts was strongly influenced by the age of fungal and mycelial vitality. This study evaluated the production of protoplasts from liquid cultured mycelia harvested at different incubation times (4, 5, 6, and 7 days). The highest protoplast yield was obtained from 4-day-old mycelia, reaching a concentration of 4.03 × 10^7^ CFU/mL (Fig. 2A). Prolonged cultivation resulted in a sharp decline in protoplast production, the yields of 5-, 6-, and 7-day-old mycelia were 2.81 × 10^7^ CFU/mL, 1.42 × 10^7^ CFU/mL, and 0.85 × 10^7^ CFU/mL, respectively (p < 0.001). This reduction suggests that prolonged culture durations lead to thickened cell walls, which hinder effective enzymatic digestion. Based on these results, 4-day-old mycelia were selected for all subsequent protoplast preparations.

**FIG 2.**
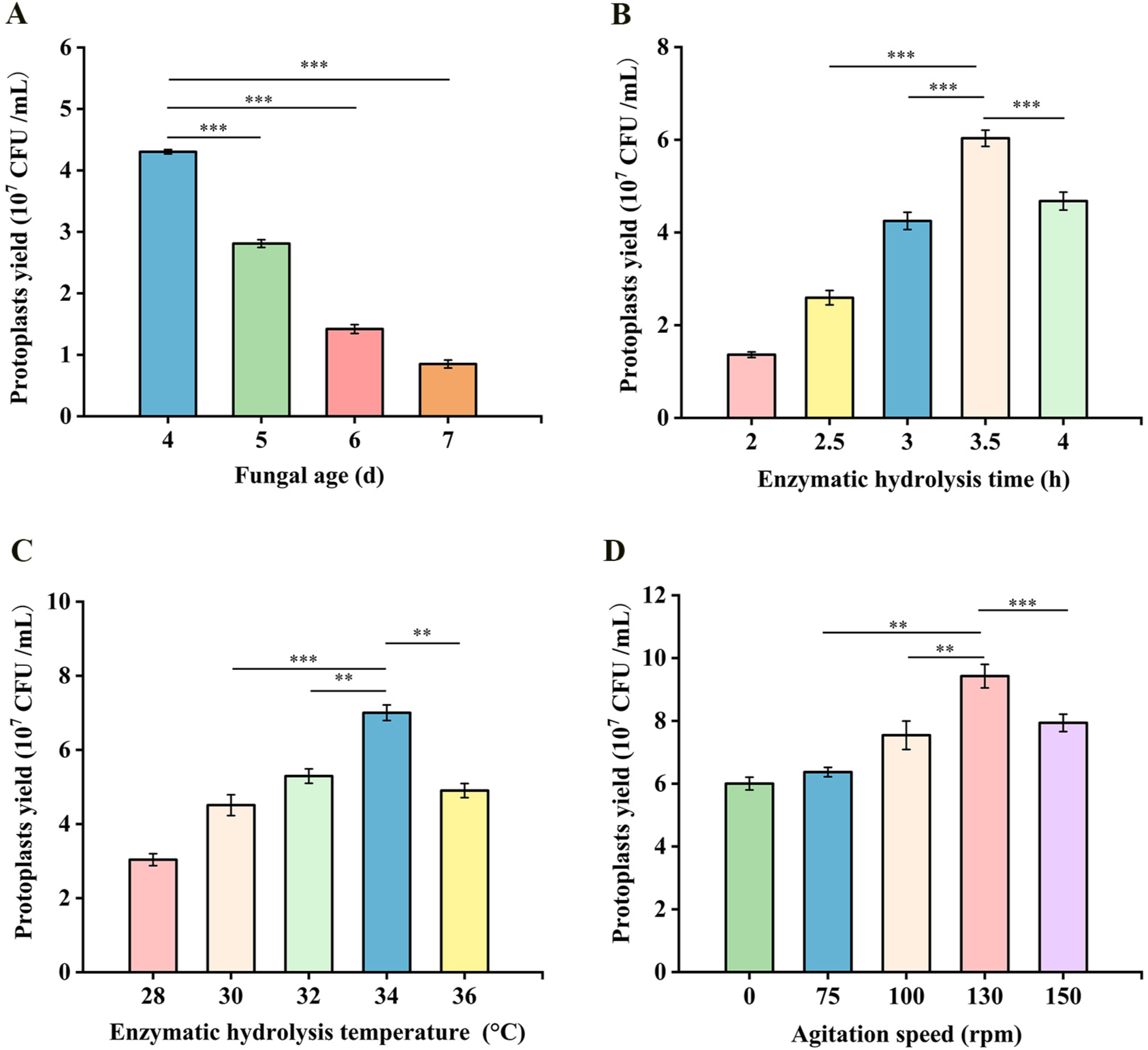
Optimization of the preparation system for *O. xuefengensis* protoplasts. (A) fungal age; (B) enzymatic hydrolysis time; (C) enzyme hydrolysis temperature; (D) agitation speed. Values are presented as mean ± SD (*n* = 3). Significance analysis between two different groups: ^∗^ *p <* 0.05, ^∗∗^ *p <* 0.01, and ^∗∗∗^ *p <* 0.001.

#### Effect of enzymatic hydrolysis environments on protoplast preparation

In addition to fungal age, enzyme composition, and concentration, environmental factors during enzymatic digestion, such as the enzymatic digestion time, enzymatic digestion temperature, and agitation speed, also have a significant impact on protoplast yield. The number of *O. xuefengensis* protoplast increased rapidly with increasing enzymatic digestion time, reaching a maximum yield of 6.03 × 10^7^ CFU/mL after 3.5 h (Fig. 2B). However, when digestion was extended to 4 h, the protoplast yield decreased to 4.68 × 10^7^ CFU/mL (p < 0.001), indicating that prolonged enzymatic digestion leads to protoplast rupture. Therefore, an enzymatic hydrolysis time of 3.5 h was selected as the preferred time for protoplast preparation.

The protoplast yield exhibited a positive correlation with the enzymatic digestion temperature in the range of 28–36^◦^C (Fig. 2C). The highest yield, 7.01 × 10^7^ CFU/mL, was achieved at 34^◦^C following cell wall digestion, significantly surpassing those obtained at 28^◦^C (3.03 × 10^7^ CFU/mL, p < 0.001), 30^◦^C (4.51 × 10^7^ CFU/mL, p < 0.001), and 32^◦^C (5.29 × 10^7^ CFU/mL, p < 0.01). Beyond this temperature, protoplast production decreased, reaching 4.90 × 10^7^ CFU/mL at 36^◦^C. Based on these results, 34^◦^C was therefore identified as the optimal temperature for subsequent experimental procedures.

Furthermore, the effect of agitation speed on protoplast preparation was evaluated, four rotational speeds (75, 100, 130, and 150 rpm) were applied while maintaining previously optimized enzymatic hydrolysis conditions (Fig. 2D). The protoplast yields were significantly higher under all agitated conditions than those under static digestion. The highest protoplast yield of 9.42 × 10^7^ CFU/mL was obtained at 130 rpm. Deviation from this speed resulted in reduced protoplast production: yields of 7.54 × 10^7^ CFU/mL and 7.93 × 10^7^ CFU/mL (p < 0.01) of protoplasts were obtained at 100 rpm and 150 rpm, respectively. In summary, the optimal protoplast preparation protocol consisted of digesting 4-day-old mycelia for 3.5 h at 34^◦^C with agitation at 130 rpm using an enzyme mixture of 1.5% lywallzyme and 1.5% snailase. These conditions provide a reliable foundation for subsequent transformation experiments.

### Protoplast regeneration

The obtained protoplasts were diluted to a concentration of 1 × 10^7^ CFU/mL. 0.6 M sorbitol was selected as the osmotic pressure stabilizer, three regeneration media (PPDA, PY and TB3) were evaluated to determine the regeneration efficiency of *O. xuefengensis* protoplasts and identify the most suitable regeneration medium. Regeneration of the protoplast occurred successfully in all tested media (Fig. 3). In particular, visible colonies on the PPY medium almost covered the entire plate, exhibiting markedly higher regeneration compared to PPDA and TB3 medium. This result clearly indicates that the PPY medium provides a more favorable environment for protoplast recovery. On the basis of these findings, the PPY liquid low melting point medium was selected for all subsequent protoplast regeneration and transformation procedures.

**FIG 3.**
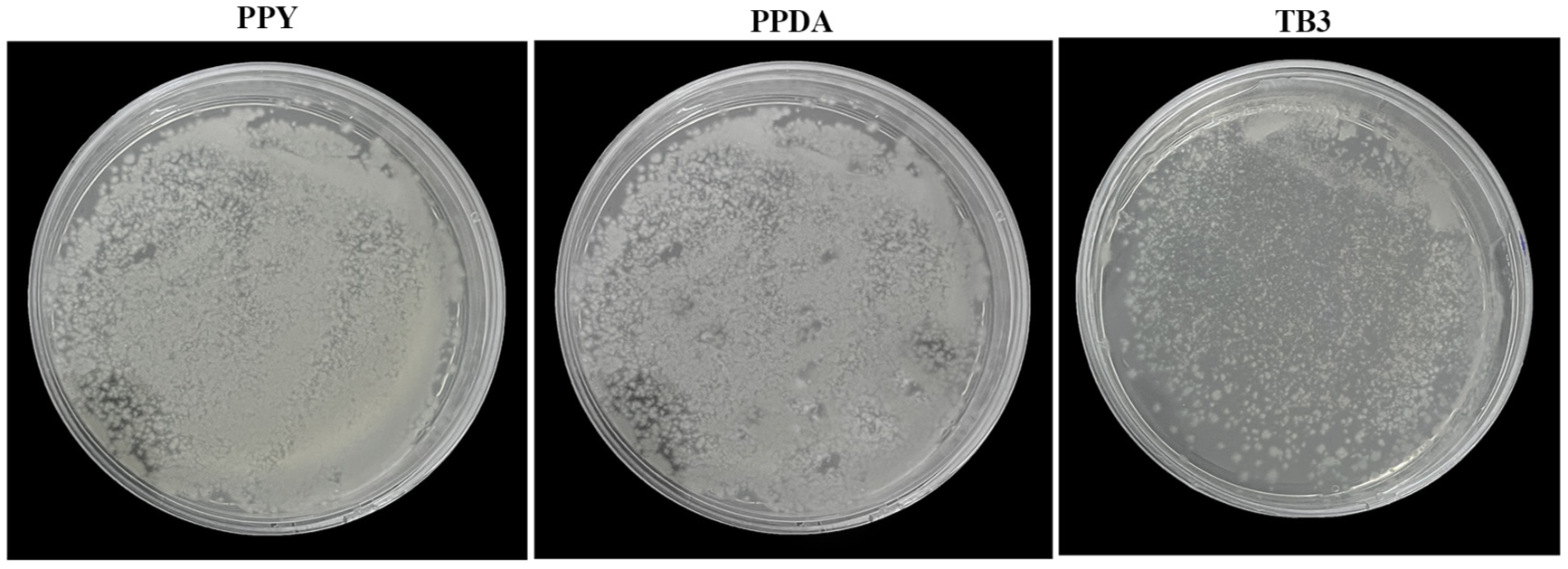
Effects of different regeneration media on *O. xuefengensis* protoplasts regeneration. PPY: PY + mannitol medium. PPDA: PDA + mannitol medium. TB3 medium.

### Resistance gene selection and selection pressure

Hygromycin B has been widely used in the genetic transformation of many fungi and is an ideal screening reagent due to its high selection efficiency and small genotype differences [33]. In this study, we systematically evaluated the sensitivity of O. xuefengensis to hygromycin B. The diameter of the plate strain gradually decreased as the concentration of hygromycin B increased from 0 to 600 µg/mL (Fig. 4). At a concentration of 600 µg/mL, no growth was observed. Therefore, 600 µg/mL of hygromycin B was selected as the preferred concentration for the subsequent screening of transformants.

**FIG 4.**
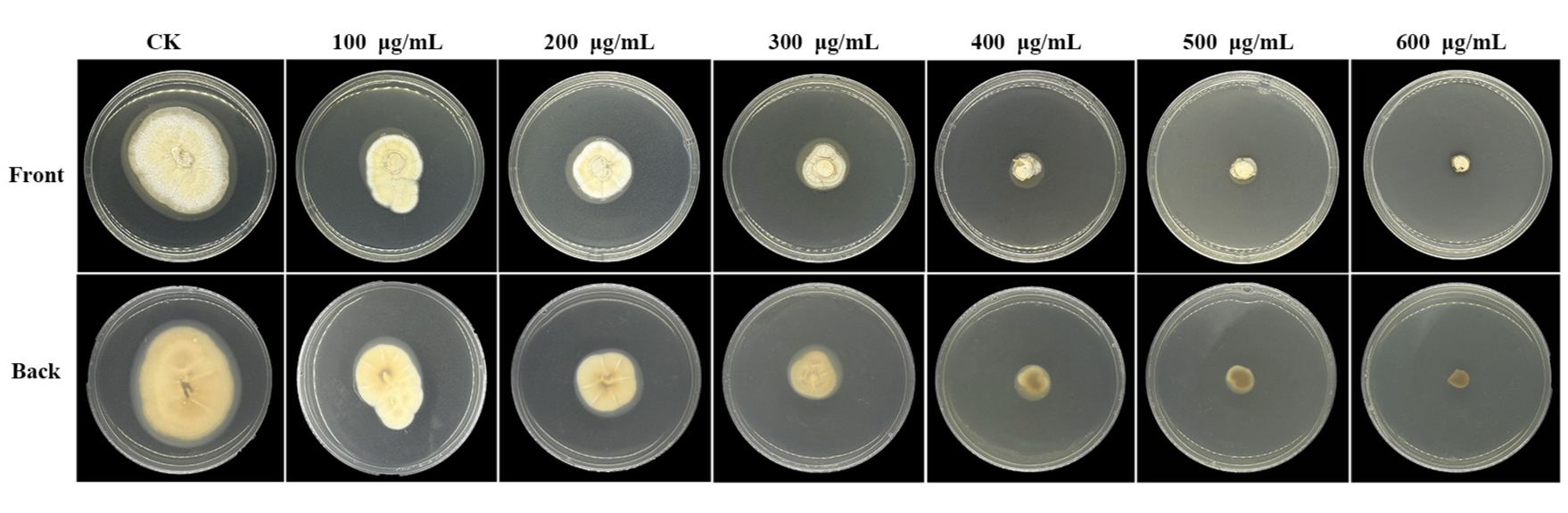
Sensitivity of *O. xuefengensis* HACM001 to hygromycin B at different concentrations of hygromycin B (200–600) µg/mL) on PY medium, wild-type strain without hygromycin B was as negative control (CK).

### PEG-Mediated transformation and verification of transformants of *O.xuefengensis*

To evaluate the transformation efficiency of *O.xuefengensis* protoplasts, the plasmid which contains hygromycin B resistance (*hygR*) gene was integrated into the *O. xuefengensis* genome using a PEG-mediated transformation approach. Compared to wild-type control, transformants carrying the *hygR* gene exhibited robust growth in PPY medium supplemented with hygromycin B (Fig. 5A). PCR amplification confirmed the presence of the *hygR* gene in the transformants, and six independent lines were obtained that showed correct integration and high genetic stability (Fig. 5B). These results confirm the successful establishment of a genetic transformation system in *O. xuefengensis*.

**FIG 5.**
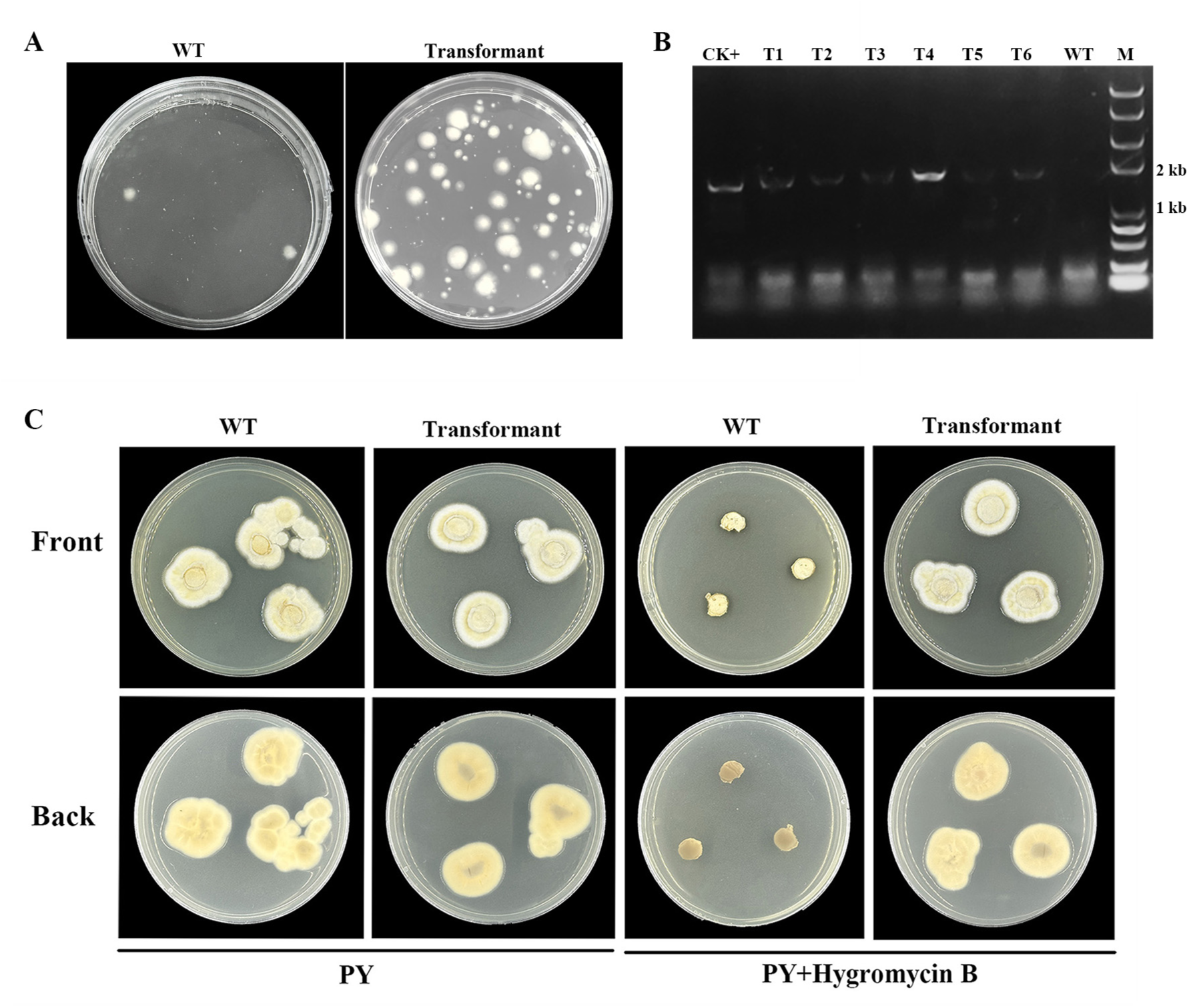
Comparison of colony regeneration before and after transformation with pCAMBIA1300, molecular verification of transformants and genetic stability analysis after four generations of cultivation. (A) Growth of untransformed (left) and transformed (right) protoplasts on hygromycin B-containing PPY medium. (B) PCR verification of the *hygR* gene in wild-type (WT) and putative transformants (T1-T6) using gene-specific primers. M: marker. CK+: positive control (plasmid pCAMBIA1300). (C) Phenotypes of wild type strain *O. xuefengensis* HACM001 (WT) and fourth positive transformants (transformant) on antibiotic medium at 28^◦^C for 10 days.

To evaluate the genetic stability of the *hygR* gene, three of the first generation transformants were randomly selected and initially cultured in hygromycin-containing PY medium.Subsequently, these were subsequently subcultured twice in antibiotic-free PY medium to assess retention of the resistance marker in the absence of selection pressure. The fourth generation was then replated on hygromycin-supplemented medium, and PCR analysis again confirmed the presence of the *hygR* gene in all transformants. The fourth positive transformants exhibited stable growth in hygromycin-containing selection medium (Fig. 5C), demonstrating that the *hygR* gene was stably integrated into the fungal genome and maintained for multiple generations under selective and non-selective conditions.

### Evaluation of PEG-Mediated protoplast transformation on heterologous CBHI expression

To further evaluate the feasibility of the polyethylene glycol (PEG)-mediated genetic transformation system, the recombinant expression cassette pCAMBIA1300-CBHI was first transformed into *O. xuefengensis* protoplasts according to the transformation method mentioned above. The pCAMBIA1300-CBHI recombinant expression cassette was successfully constructed by homologous recombination of the endogenous promoter P*_Oxgpd1_*, the exogenous glycoside hydrolase gene *cbhI* (encoding cellobiohydrolase I, CBHI), and the terminator *trpc* (Fig. 6A).

**FIG 6.**
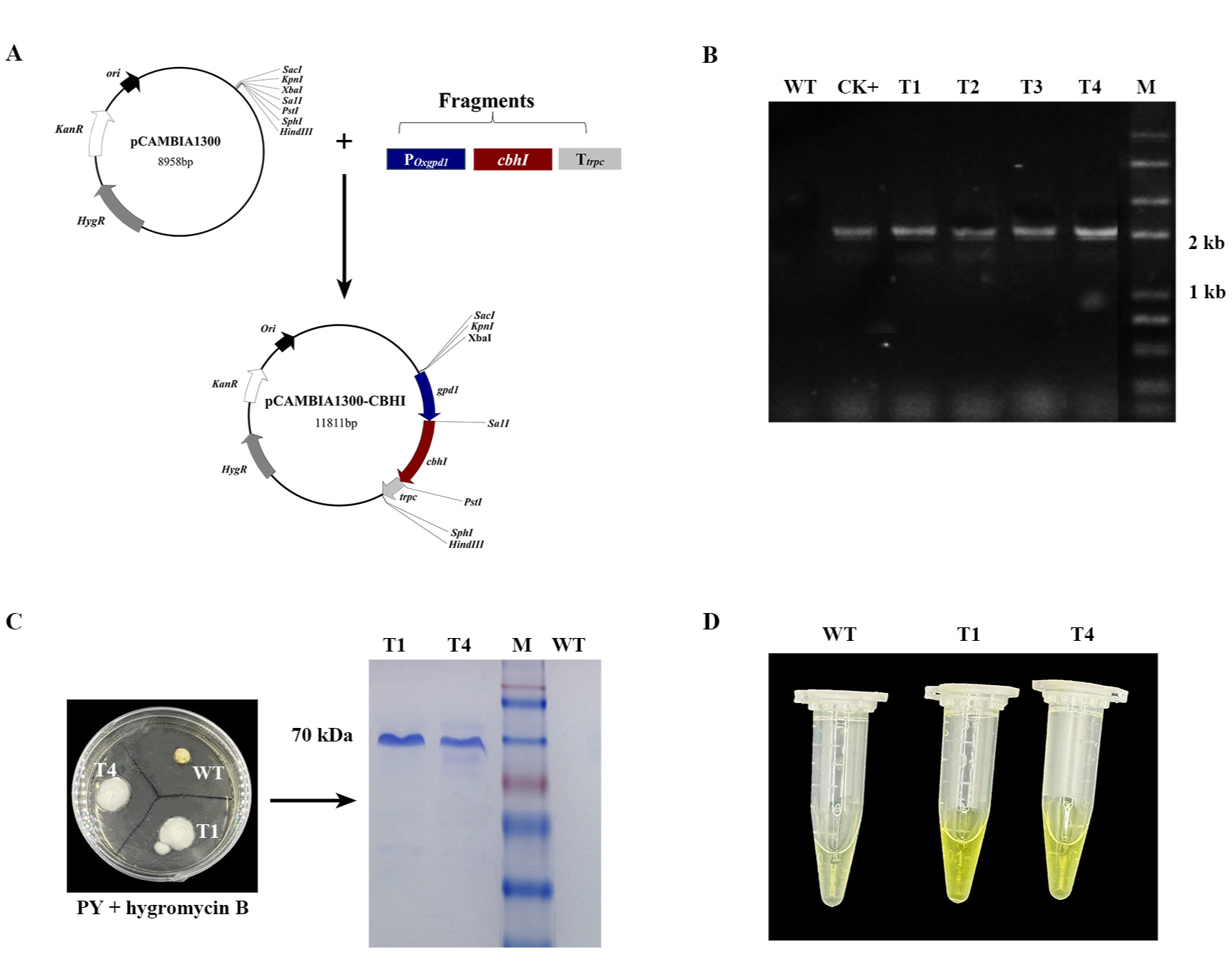
Genetic transformation feasibility of exogenous glycoside hydrolase gene *cbhI* in *O. xuefengensis*. (A) The construction of recombinant expression cassette pCAMBIA1300-CBHI. (B) PCR amplification of the *gpd1*+*cbhI* gene (2026 bp). (C) Phenotypes of wild type strain (WT) and transformants (T1 and T4) on antibiotic medium (left), and SDS-PAGE analysis of the purified CBHI protein(right). (D) Analysis of enzyme activity of the purified CBHI protein. CK+: Positive control (plasmid pCAMBIA1300-CBHI). T1-4: Transformant 1-4. M: marker.

The transformants were initially selected on PPY medium and subsequently subjected to three rounds of antibiotic screening. The successful integration of the desired gene was verified by amplifying 2.02 kb of the glycoside hydrolase gene *cbhI* into the genomic DNA of *O. xuefengensis* of randomly selected transformants, while no amplification product was detected in the wild-type strain (Fig. 6B). Furthermore, two positive transformants growing on a resistance plate were cultured in fermentation broth and extracellular proteins were extracted and purified. SDS-PAGE analysis confirmed the successful expression and secretion of CBHI (Fig. 6C). Subsequent enzymatic assays demonstrated that the recombinant CBHI exhibited cellobiose hydrolyzing activity, confirming its functional expression (Fig. 6D). These results collectively demonstrate the reliability of the established transformation system and provide a solid technical foundation for future functional genomic studies in *O. xuefengensis*, including gene knockout, overexpression, and investigation of genes regulating fruiting body development.

## DISCUSSION

*Ophiocordyceps xuefengensis* is an emerging medicinal resource known for its diverse bioactive compounds. In-depth investigation of the biosynthesis of its metabolites and the regulatory mechanisms underlying its growth and development are crucial to expanding its potential applications. However, progress in functional genes and research on secondary metabolite synthesis in *O. xuefengensis* has been hampered by the lack of an efficient platform of genetic transformation. Although genetic transformation techniques have been successfully established in other species of Cordyceps and edible fungi, transformation strategies exhibit considerable variation between taxonomic groups [34, 35]. Therefore, it is urgent to develop a stable and reproducible genetic transformation system for *O. xuefengensis* to facilitate molecular studies and breeding efforts.

The successful establishment of a genetic transformation system in pathogenic bacteria and filamentous fungi is highly dependent on the yield and quality of protoplasts [36]. The complex composition and structure of the fungal cell wall significantly influence the efficiency of protoplast preparation. Substantial evidence indicates that the type and concentration of cell wall lytic enzymes are strongly correlated with the production of protoplast [37, 38]. Appropriate and highly active enzyme mixtures can digest the mycelial cell wall more effectively, thereby releasing a greater number of intact protoplasts. In this study, six enzymatic combinations were evaluated for their efficacy in protoplast preparation from *O. xuefengensis*. Combination 4 (lywallzyme1 and snailase) released 2.23 × 10^6^ CFU/mL of protoplast after 4 h of digestion, which was significantly higher than that of other groups (Fig. 1A) and therefore was selected for subsequent experiments. This approach using multiple and mixed enzyme systems for the preparation of protoplasts is consistent with strategies applied in other fungal systems. For example, a four-enzyme mixture has been used for efficient protoplast isolation from *Penicillium sclerotiorum* mycelium, while a two-enzyme combination was shown to be optimal for cell wall digestion and protoplast release from *Colletotrichum falcatum* [39, 40]. In contrast, some species such as *cordyceps cicadae* and *Trichoderma reesei* require only a single enzyme for effective protoplast preparation [20, 41]. Furthermore, the concentration of lytic enzymes is critically associated with the yield of protoplast [42].

However, beyond a certain threshold, increasing the concentration of enzymes does not substantially improve the efficiency of release, highlighting the importance of identifying optimal digestion conditions [43]. Consistent with this principle, our results demonstrated that a combination of 1.5% lywallzyme 1 and 1.5% snailase significantly enhanced the protoplast yield, reaching 9.11 × 10^6^ CFU/mL (Fig. 1B).

The age of the mycelium, along with the enzymatic digestion environments, including temperature, time, and agitation speed, are critical factors affecting the yield and viability of protoplasts [44, 45]. The structure and metabolic activity of the cell wall vary considerably between developmental phases. Very young mycelia exhibit thin and fragile cell walls, whereas older mycelia develop thick and rigid walls, both states are suboptimal for protoplast isolation [46]. Previous studies indicate that young, actively growing mycelia provide the best balance between structural integrity and digestibility, making them the most suitable for efficient protoplast preparation [47]. Furthermore, insufficient digestion time can result in incomplete cell wall lysis, while excessive duration can cause protoplast rupture and diminish regeneration capacity [20]. The temperature of digestion directly influences the enzyme kinetics, and moderate agitation enhances enzyme-substrate interaction and digestion homogeneity [48]. In this study, the highest yield of protoplasts (9.42 × 10^7^ CFU/mL) was obtained using 4-day-old mycelia digested at 34^◦^C with agitation at 130 rpm for 3.5 h with 1.5% lywallzyme 1 and 1.5% snailase (Fig. 2D). Regeneration of the protoplast is equally critical as preparation and requires an optimized regeneration medium. The choice of osmotic stabilizers varies between fungal species, for example, mannitol is used for *Cordyceps militaris* [49], KCl for *Ganoderma lucidum* [50], sorbitol for *penicillium oxalicum* [16]. In this study, PY medium supplemented with 0.6 M mannitol supported the highest regeneration rate for *O. xuefengensis* protoplasts (Fig. 3), consistent with previous reports that appropriate osmotic stabilizers help preserve protoplast integrity and promote cell wall regeneration [51, 52].

The selection of a suitable selectable marker and its corresponding inhibitory concentration is crucial to a successful fungal transformation [53]. Hygromycin B is widely used to screen transformants in ascomycetous fungi, with effective concentrations ranging typically from 50 to 400 µg/mL [37, 54]. In this study, the minimum inhibitory concentration (MIC) of hygromycin B for *O. xuefengensis* was determined to be 600 µg/mL (Fig. 4), indicating a relatively high resistance compared to other fungi. This finding is consistent with previous reports by Sun et al., who employed 650 mg/L hygromycin for selecting transformants of *Cordyceps militaris* [55]. For PEG-mediated transformation, concentrations of PEG 4000 between 200 and 400 g/L are commonly employed [56, 57].

Using 250 g/L PEG 4000, we successfully achieved genetic transformation in *O. xuefengensis* first time, as confirmed by antibiotic selection and PCR verification of positive transformants. The transformation efficiency was approximately 50 transformants per 6 µg of plasmid DNA, with the transformants obtained showing stable genetic inheritance through successive generations (Fig. 5C).

Endogenous promoters are more readily recognized by the transcriptional machinery of the host organism, significantly improving both genomic integration efficiency and transcription initiation [27, 58]. To further evaluate the feasibility of the transformation system, we employed the endogenous GPD promoter from *O. xuefengensis* (P*_Oxgpd1_*) to drive the expression of the *cbhI* gene. The recombinant vector pCAMB1A300-CBHI was successfully integrated into the fungal genome, and the purified enzyme of CBHI demonstrated the specific hydrolyzing activity of cellobiose (Fig. 6D), which confirms the functional expression of the heterologous gene. The genetic transformation system established in this study provides a robust platform for functional genomics research in *O. xuefengensis*. Future studies should focus on elucidating the molecular mechanisms underlying growth and development in this species to facilitate the breeding of improved strains with enhanced medicinal properties.

## ACKNOWLEDGMENTS

The author thanks Liping Lin for her assistance in optimizing protoplast preparation and Yifan Jiang for her assistance in the CBHI enzyme activity determination experiment at Changsha University of Science and Technology.

This article was supported by the Natural Science Foundation of Hunan Province of China (No. 2023JJ40017, 2025JJ60526, 2024JJ7292), the Health Research Project of Hunan Provincial Health Commission (No. 20255256), the Key project at central government level: The ability establishment of sustainable use for valuable Chinese medicine resources (No. 2060302-2304-02, 2060302-2503-22), and Postgraduate Scientific Research Innovation Project of Hunan University of Chinese Medicine (No. 2025CX187).

## FUNDING

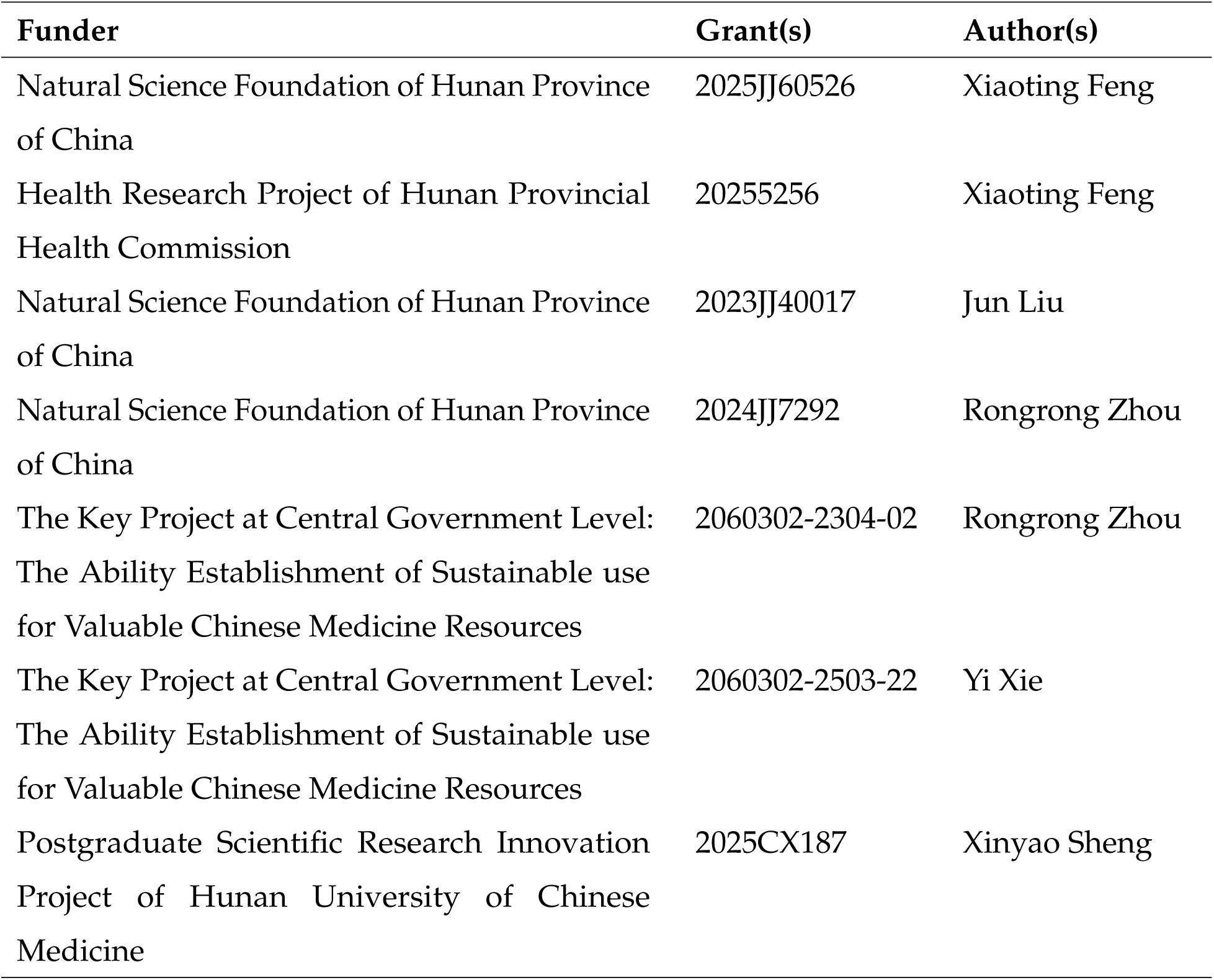

## AUTHOR CONTRIBUTIONS

XF: Conceptualization, Formal analysis, Investigation, Data curation, Writing – original draft, Writing-review & editing, Validation, and funding Acquisition. XS: Formal analysis, Data curation, Visualization. JL and RZ: Methodology, Writing-review & editing and Funding acquisition. ZY and XT: Formal analysis, Investigation. SZ: Conceptualization, Funding acquisition, Project administration, Supervision.

